# Urbanization drives the acoustic diversity of mangrove anurans and avifauna in Puerto Rico

**DOI:** 10.1101/506139

**Authors:** Benjamin Branoff

## Abstract

The mangroves of Puerto Rico occupy a broad gradient of urbanization that offers a chance to test hypotheses on urban faunal communities from both terrestrial and mangrove systems. These hypotheses state that urban avifaunal communities are less diverse, with greater representation by generalists, but that certain mangrove specialists can utilize urban landscapes. Much of this is said to be driven by food resources, with frugivores and nectarivores benefiting from abundant residential flowers and fruits, while insectivores are driven away by low food resources. This study used passive acoustic monitoring to identify the audible anuran and avifaunal species in mangroves across the urban gradient of Puerto Rico. Three anurans and twenty-four avian species were detected across all sites, with eighteen species found at the most diverse site, and six at the least diverse site. The most urban communities were more similar to each other and characterized by low diversity, especially of invertivores, mangrove specialists, and endemics. The least urban communities, however, were less like each other and more diverse than their more urban counterparts. Greater surrounding vegetation cover was the strongest overall predictor of increased faunal diversity, although urban coverage and population density were also strongly correlated with some guilds and species. Soundscape analyses showed that acoustic space use was a strong predictor of site diversity and is thus a sufficiently accurate faster alternative to detecting individual species. This will be an important tool in the continued monitoring of these forests, which may experience changes associated with ongoing urbanization and sea-level rise.

## Introduction

Urban bird communities have repeatedly been shown to change in composition with increasing urbanization (Chace and Walsh, 2006; Clergeau et al., 2001, 1998; Emlen, 1974; Ortega-Álvarez and MacGregor-Fors, 2009; Tilghman, 1987). Generally urban avifauna are mostly represented by generalist species capable of foraging on a broad range of food types, with the more specialist species largely absent from the most urban sites (Chace and Walsh, 2006). Most of these studies also acknowledge the importance of various metrics of the surrounding land cover matrix, such as forest patch size or habitat diversity. These patterns cover a broad range of habitats and latitudes, and similar observations have been made concerning the urban avifauna of mangroves.

A number of studies have examined urban mangrove avifaunal communities (Acevedo and Aide, 2008; Lim and Sodhi, 2004; Mancini et al., 2018; Mestre et al., 2007; Mohd-Azlan and Lawes, 2011; Pawar, 2011; Ward, 1968), but there is no agreement between them on what constitutes “urban” and few conclude on the overall influence of urbanization on community structure, with some exceptions (Lim and Sodhi, 2004; Noske, 1996, 1995; Ward, 1968). Several studies acknowledge a relatively high presence of mangrove specialists in anthropogenic habitats of Malaysia (Lim and Sodhi, 2004; Noske, 1995; Ward, 1968), suggesting these species are particularly adept at colonizing urban landscapes. But another study in Australia shows that most of the mangrove specialists there have yet to colonize the urban mangroves of Darwin (Noske, 1996). A different study in Darwin suggests the richness of the mangrove bird community is most dependent upon the variation in the surrounding land use matrix (Mohd-Azlan and Lawes, 2011). In Puerto Rico, apart from studies on specific species or generalized ecosystem studies, only one study has focused on the mangrove avifauna of the island (Acevedo and Aide, 2008), and none have studied its mangrove anurans.

Acevedo and Aide (2008) found twenty-six species of birds in one mangrove forest of Puerto Rico, with significant differences in community assemblages during the migratory and non-migratory seasons. While the detection of resident species did not change from season to season, the detection of migrants did, increasing during the migratory season as expected. The authors suggest that these migrants are more strongly represented in the mangroves in comparison to other habitats due to their primarily insectivorous diet and an abundance of insect food resources in wetland habitats. This study also notes a relationship between the mean tree size in the forests and the associated bird community. Another survey conducted mostly in, but not limited to the mangroves of San Juan, found a maximum site richness of thirty-one aquatic species, with both richness and abundance positively proportional to the surrounding area of mangroves (Fidalgo-De Souza, 2009). Outside of mangroves, one study showed certain generalist anuran and avian species are more associated with urban landscapes than with reference forested sites on the island (Ruiz-Jaén and Aide, 2006), while another has shown how human activity and acoustic sound influence avian and anuran communities (Herrera-Montes and Aide, 2011).

Some of the above studies utilize bioacoustic surveys to quantify species presence. This method has repeatedly been identified as being similar to or more effective than visual point counts in a variety of ecosystems (Alquezar and Machado, 2015; Haselmayer and Quinn, 2000; Zwart et al., 2014), and especially in mangroves (Celis-Murillo et al., 2012). The primary limitation to this method is the need for large amounts of digital storage and the development of time demanding acoustic pattern recognition models. Aide et al. (2017) suggest a potential alternative to these models for generalizing the relative diversity between sites: Acoustic Space Use (ASU). ASU is the percentage of acoustic space, comprised of both time and frequency ranges, used by the collective sound of the faunal community. In their study, Aide et al. (2017) show that ASU at a site is directly related to its faunal diversity, especially of insects. Thus, ASU is relatively easily calculated metric of biodiversity that could be used for rapid assessments and long-term monitoring of noisy faunal communities.

This study uses acoustic surveys and subsequent acoustic pattern recognition models to detect avian and anuran presence in twenty forested mangrove sites spanning an urbanization gradient in Puerto Rico. It then aims to answer how mangrove avian and anuran diversity is related to various metrics of surrounding urbanization and land cover, focusing on specific feeding guilds, endemics, and mangrove specialists, as well overall diversity and the abundance of individual species’ vocalizations. It then shows how a site’s diversity is related to its ASU, and the role of anthrophony in this relationship. This is a comprehensive account of the avian and anuran communities of Puerto Rico’s mangroves, and the presented relationships can be used to guide conservation and management of these highly social-ecological systems.

## Methods

### Site Description

The urban mangroves of Puerto Rico, and the sites corresponding to the present study, are described in detail in Branoff and Martinuzzi (2018). Surface water chemistry and flooding dynamics are described in (Branoff, 2018). Twenty one-hectare forested areas were selected among three watersheds, which were chosen because they harbored mangroves with the greatest range in urbanization (Figure 1). Urbanization was determined by sampling surrounding spatial datasets of urban land cover (impervious surfaces), vegetation cover, mangrove cover, road density, and population density, as described by (Branoff and Martinuzzi, 2018). The three watersheds are described as the Río Hondo to the Río Puerto Nuevo, referred to here as San Juan; the Río de la Plata, referred to here as Levittown; and the Río Inabón to the Río Loco, referred to here as Ponce. The twenty sites were chosen to span the greatest range in urbanization within the watersheds. Fourteen sites were selected in San Juan, two in each waterbody representing the minimum and maximum urban sites. Three sites were selected in each of the Levittown and Ponce watersheds. Site abbreviations for Levittown and Ponce begin with LEV and PON, respectively, while those in San Juan begin with the waterbody name: BAH for Bahía de San Juan, MPN for Martin Peña non-dredged, MPD for Martin Peña dredged, PIN for Piñones lagoon, SAN for San José lagoon, SUA for Suarez Canal, and TOR for Torecillas lagoon. These are followed by MIN, MID, or MAX referring to the relative level of urbanization in each waterbody or watershed. Five sites in San Juan lie within 0.5 km of low-altitude commercial airline flight paths, which subjects these sites to periodic high intensity sound. These are MPDMIN, MPDMAX, MPNMIN, MPNMAX and TORMIN.

**Figure 1:**
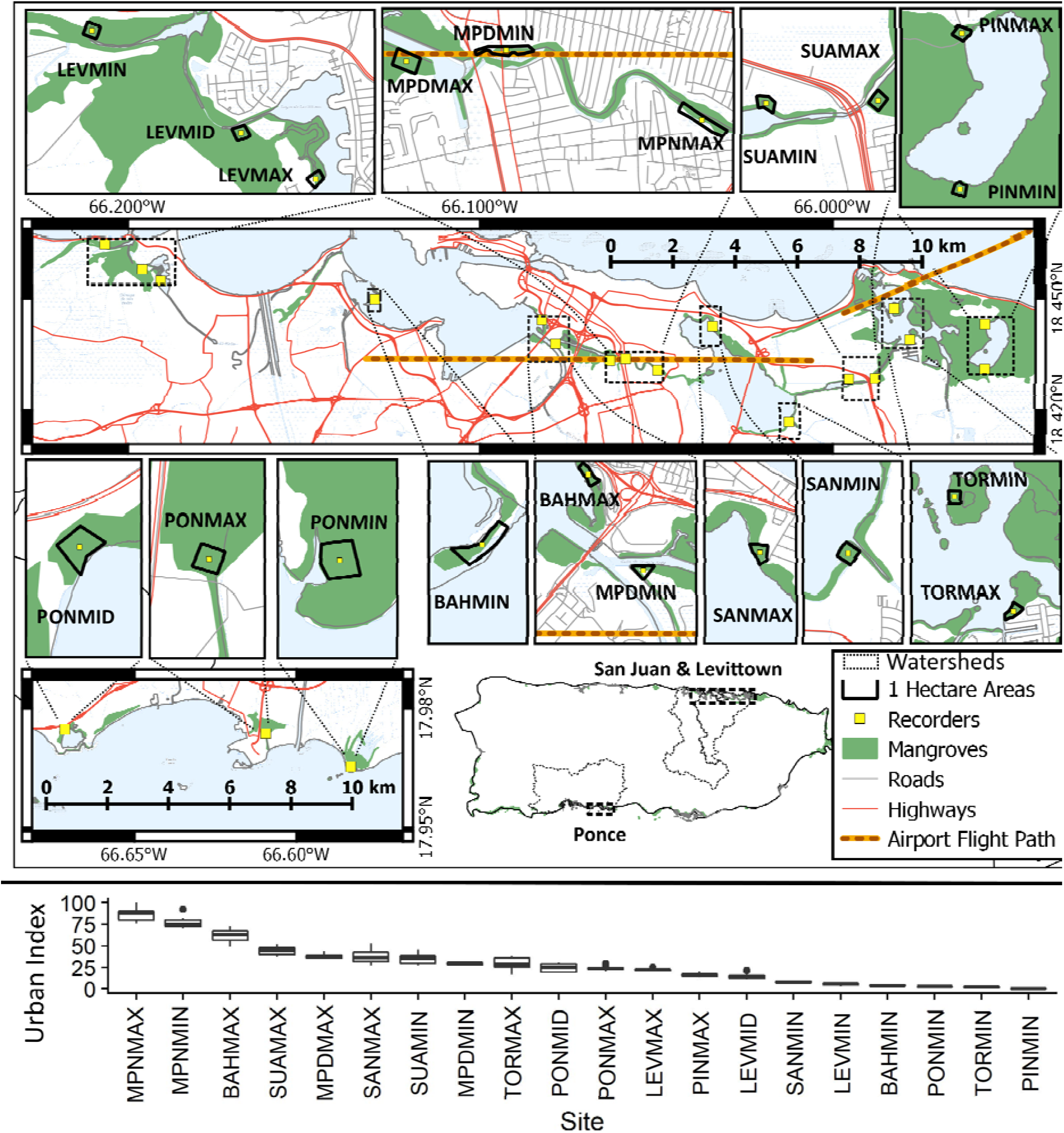
Passive acoustic monitoring devices were placed in twenty one-hectare mangrove forests spanning an urbanization gradient across the island of Puerto Rico, as represented by the urban index. This index is calculated using the surrounding cover of urban, vegetated, open water, and mangrove classifications, as well as road and population density. Higher values are the most urban sites.

Forest composition and structure vary between sites, but in general the forests are composed primarily of *Laguncularia racemosa*, followed by *Rhizophora mangle* and *Avicennia germinans*. Additional, non-halophytic flora are present, and the overall floral diversity increases with population density surrounding each forest (Branoff and Martinuzzi, 2018). The diversity of the three mangrove species, however, decreases as urban cover increases. Forest height varies between ten and twenty meters, stem density between 5 and 7.7 thousand stems per hectare, and basal area between 27 and 43 m^2^ per hectare. Most of the structural metrics are correlated with surrounding vegetation cover or surface water chemistry, with the largest contiguous forests being characterized as taller and with greater variation in height.

Relatively little is known of the mangrove fauna of Puerto Rico in comparison to other ecosystems on the island. The most urban site at Caño Martín Peña (MPNMAX) was surveyed in 1899 and noted the presence of a diverse avian community composed in part of *Megaceryle alcyon, Pelecanus occidentalis, Ardea spp*., *Rallus sp.*, Scolopacidae sp., *Setophaga spp*., *Tyrannus dominicensis* and *Vireo altiloquus* (Evermann et al., 1902). Another more recent survey found the aquatic bird community in San Juan to be composed mostly of *Fulica caribea, Bubulcus ibis, Pluvialis squatarola, Pelecanus occidentalis*, and *Calidris alba*, although this study was not limited to mangroves (Fidalgo-De Souza, 2009). Acevedo and Aide (2008) surveyed the bird community of the Levittown mangroves and found twenty-six species, with *Coereba flaveola, Quiscalus niger, Seirus noveboracensis*, and *Melanerpes portoricensis* to be among the most abundant. Descriptions of the mangrove bird community in Ponce are absent, but other studies along the southern coast of the island note the use of mangroves for both foraging and nighttime roosting habitat, with the most abundant species being *Setophaga petechia* and *Seiurus noveboracensis* (Rodríguez-Colón, 2012; Smith et al., 2008).

### Faunal Surveys

Two surveys were conducted in each site, one from March 11^th^ to March 22^nd^, and from November 1^st^ to November 12^th^, 2017. Passive acoustic monitoring devices were fixed to a tree at a height of approximately 1.5 meters at each site and programmed to sample for 1 min at 10-min intervals at a sampling rate of 44.1 kHz. Recorders are comprised of an Android smartphone attached to an external Monoprice microphone with flat response between 50 Hz and 20 kHz and a sensitivity of −45 ± 2 dB, all enclosed in a waterproof case. Recorders were programmed with the ARBIMON Touch application (https://goo.gl/CbBavY) and allowed to record until the smartphone battery died, or until they were collected ten days later. All recordings were stored, processed, and analyzed using the ARBIMON II platform (https://arbimon.sieve-analytics.com/)(Aide et al., 2013).

All recordings were visually inspected for interference or damage caused by temporary microphone malfunction and these recording were removed form the analyses. Species specific acoustic models were constructed by first finding the most representative, loud, and clear examples of each species’ most consistent vocalization. Species identifications were checked with those found on xeno-canto.com and by an expert in Puerto Rican avian and anuran fauna. In some cases, similar species could not be differentiated and are referred to by their lowest confirmed taxonomic rank. Representative examples of each species’ vocalizations, along with validated presence and absence recordings were used to train Hidden Markov Models to recognize the selected acoustic patterns (Aide et al., 2013). Models were considered sufficiently accurate if they achieved at least 80% accuracy and precision in at least ten of the validated recordings. These models were then used to scan either day or nighttime recordings, depending upon the specie’s behavior, for the presence of each vocalization. All recordings marked as present for a given species were then visually scanned by the author to verify its presence. Additionally, every tenth recording marked as absent was visually scanned to verify a specie’s absence.

To calculate ASU, soundscapes of each site were constructed within the ARBIMON platform. Recordings at each site were aggregated by the hour, with a frequency bin size of 86 Hz and an amplitude filtering threshold of 0.04. This resulted in a matrix for each site composed of the time of day as the column (one for each hour), the sound frequency range in bins as the rows (one for each 86 Hz bin), and the proportion of all recordings at the site with a sound greater than the 0.04 amplitude threshold in each time-frequency bin. For this study, the maximum frequency was set to 10 kHz as there were no detected vocalizations with a higher frequency. The ASU was calculated as the percentage of the soundscape matrix with a value greater than 0, or the percentage of the soundscape with at least one occurrence of a sound greater than the 0.04 absolute amplitude.

Avian species were classified into feeding guilds of granivores, omnivores, invertivores, frugivores & nectivores, and carnivores, piscivores & scavengers following (Wilman et al., 2014). Avian endemism was determined from (Lepage, 2018) and mangrove specialization was determined through Luther and Greenberg (2009). Anuran diets and endemism were determined from Crother (1999).

All statistical tests were performed in the R language (Yan et al., 2011). Relationships between faunal diversity and detections, and predictors of land cover, forest structure and composition, and flooding were tested through linear models of the form y ∼x and y ∼ln(x) in the *lm* function. To search for grouping in the avian and anuran communities among the most and least urban mangroves, sites were first divided into those with an urban index lower than the median, referred to as least urban, and those with an urban index higher than the median, referred to as most urban. Nonmetric multidimensional scaling (NMDS) using the Bray-Curtis dissimilarity index (Bray and Curtis, 1957) was then performed on the proportion of recordings in which each species was detected at each site, as recommend by (Minchin, 1987) and as performed through functions of the vegan package (Oksanen et al., 2018). The *metaMDS* function iterates the NMDS process from random starting points and chooses the solution with the overall lowest stress and thus the most accurate representation of the sites in the ordination space (Clarke and Ainsworth, 1993). To show how urbanization correlates with the grouping of the faunal communities, one environmental vector representing the urban index of the sites was fit to the ordination through the *envfit* function. The ordination was then rotated using the *rotateMDS* function so that the horizontal axis of the ordination was parallel to the urban index vector. This allows for easier interpretation without changing the ordination space. Ordination ellipses representing the standard error of the group centroid locations were drawn for the least urban and most urban groups using the *ordiellipse* function, also from vegan.

Species accumulation curves for each site and each watershed were constructed by counting the number of time-ordered recordings necessary to detect a new species. This was done for the total number of species as well as for the percentage of the maximum number of species detected at each site or watershed. To control for the different number of sites in each watershed, the process was repeated for incremental increases in the number of sites in each watershed. Logarithmic curves were then fit to these species-accumulation plots (Ugland et al., 2003), and the number of recordings required to reach a given number of species, as well as the number of species predicted for a given sample area were calculated through the *predict* function.

## Results

Three anuran species from two families and twenty-four bird species from fifteen families were detected across all sites during the study (Table 1). Five of these species are endemic to Puerto Rico and three are introduced, the rest being native and non-endemic. Another five species have been labeled as mangrove specialists, two of which are endemic to mangroves, and the other three are dependent on mangroves. Passeriformes was the most represented order, with eleven species, followed by Anura with three species, Guiformes and Pelecaniformes each with two species, and finally Accipitriformes, Columbiformes, Coraciiformes, Cuculiformes, Galliformes, Piviformes, Psittaciformes, and Strigiformes all with one species each. Half of the detected anuran and avian species are invertivores, followed by 31% as omnivores, 15% as piscivores or carnivores, and 7% as frugivores or nectarivores. Sufficiently accurate acoustic recognition models, with an accuracy and precision greater than 80%, were developed for eighteen of these species and used to survey all recordings across all sites. From these surveys, *Eleutherodactylus coqui* had the highest mean detection rate at 36% of nighttime recordings, followed by *Zenaida spp.* at 33% and *Coereba flaveola* at 14% of daytime recordings. All other species were detected in less than 10% of either daytime or nighttime recordings.

**Table 1:**
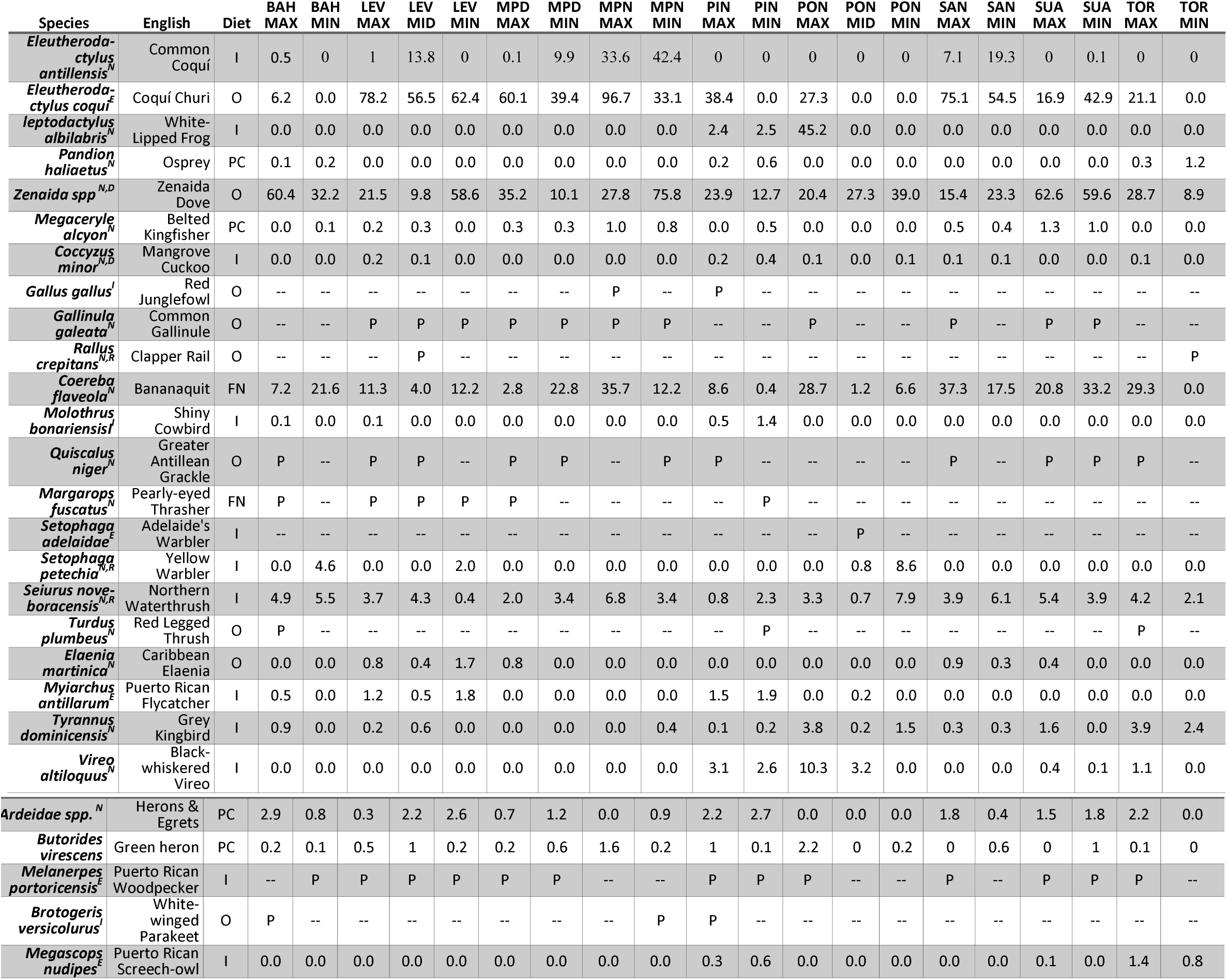
Percent species detections of all surveyed recordings at all sites for species in which acoustic pattern models were trained. Species for which models could not be constructed but were detected during non-systematic human surveys of the recordings are marked with P. Diets are Invertivore (I), Omnivore (O), Piscivore/Carnivore (PC), and Frugivore/Nectarivore (FN). Superscripts on species are native (N), endemic (E), introduced (I), mangrove dependent (D), and mangrove restricted (R). Percent species detections of all surveyed recordings at all sites for species in which acoustic pattern models were trained. P’s are species for which models ould not be constructed but were detected during non-systematic human surveys of the recordings. Diets are Invertivore (I), Omnivore (O), Piscivore/Carnivore (PC), and Frugivore/Nectarivore (FN)

The highest diversity of detected species was recorded at PINMAX, where fourteen of the modeled species and eighteen of all species were detected. Similarly, the sites of PINMIN, LEVMAX and LEVMID held seventeen of all the species, and fourteen, thirteen, and twelve of the modeled species, 1 respectively. In contrast, TORMIN held only five of the modeled species and six of all species and was the least diverse site. The site of PONMIN, PONMID, MPNMAX, and BAHMIN were similarly less diverse, with less than ten of the species recorded at all other sites. These trends were reflected in endemic diversity as well, with PINMAX containing the highest number of endemics, three, and BAHMIN and PONMIN having no recorded endemic species. Introduced species were recorded in only four of the twenty sites, at PINMAX, PINMIN, BAHMAX, and LEVMAX. There was less variation in feeding guild diversity, with all sites harboring at least one representative of all four feeding guilds, except PONMID and TORMIN, which held three guilds. In mangrove specialists, PONMIN harbored four of the five specialists detected throughout the study, while PONMID had only one.

There was a significant difference in the number of species detected at each site between the two sampling periods in March and November (t-test, mean difference = 2.4, p < 0.001). Overall, fifteen species were detected in November, compared to eighteen in March. The missing species in November were all resident species: *Setophaga petechia, Molothrus bonariensis* and *Vireo altiloquus*. Additionally, *Coereba flaveola* dropped in detection frequency across all sites by a mean of 25% from March to November. Species accumulation curves show that the mean number of minutes required to reach 50%, 75%, and 100% of all the species detected at a site are 22, 140, and 904, respectively (Figure 2). But these times varied greatly between sites. Extrapolating the species accumulation curves to 10,000 minutes for each watershed predicts 28, 24, and 21 species detected in San Juan, Levittown and Ponce, respectively. Accounting for the sampling area of each watershed brings the total predicted number of species within 15 hectares to be 28, 25, and 24 for San Juan, Levittown, and Ponce, respectively.

**Figure 2:**
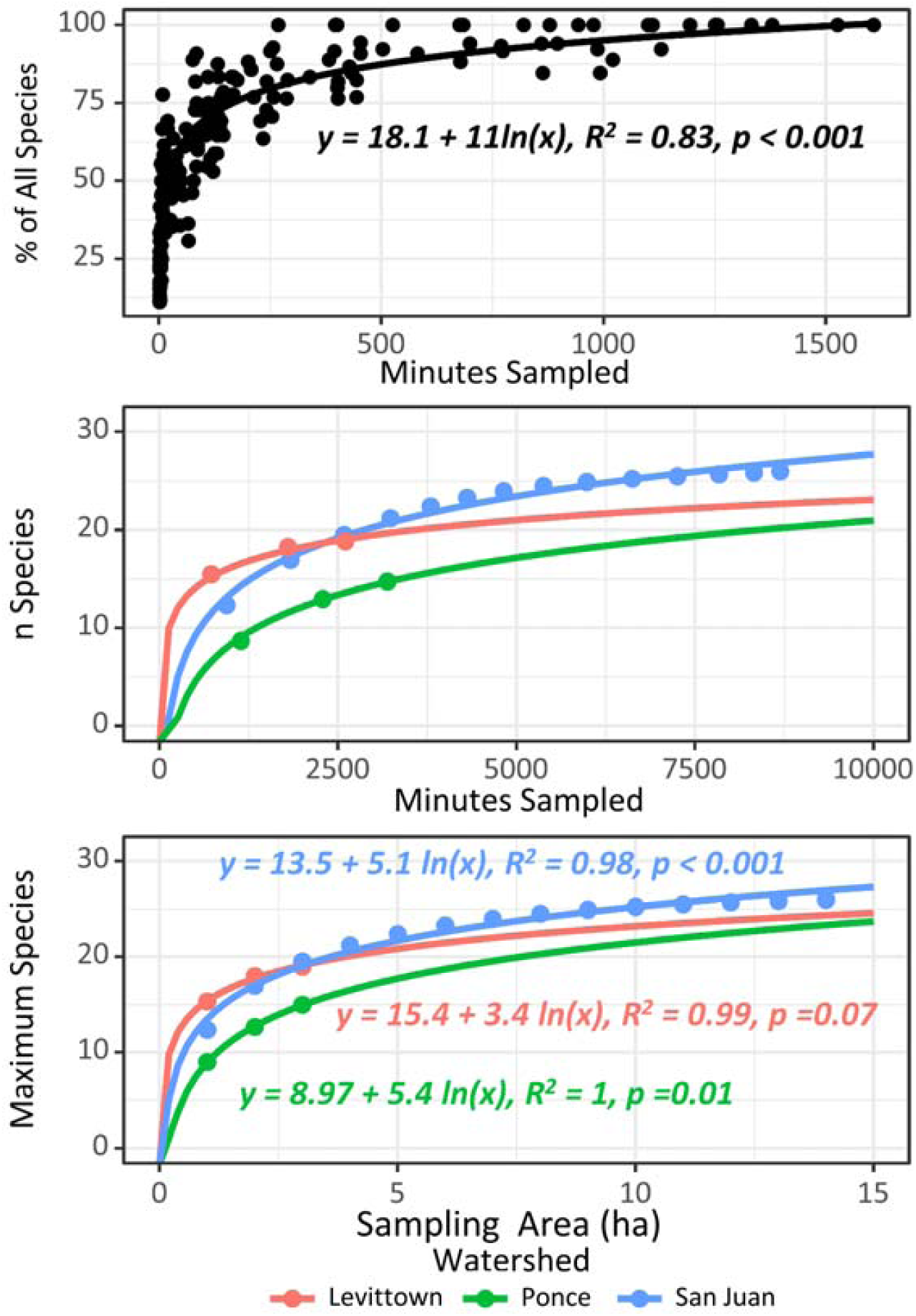
Species accumulation curves suggest an average of 22, 140, and 904 recordings to reach 50%, 75% and 100% of detected species at each site. When extrapolated, all three watersheds are predicted to harbor similar diversity, twenty-five to thirty species in 15 sampled hectares.

Differences in diversity across all sites were most strongly correlated with surrounding land cover (Figure 3). The total number of detected species at any site was positively and linearly related to the surrounding vegetation cover within one kilometer, which was also true for the number of endemic species at a site. Within feeding guilds, the invertivore diversity decreased strongly with the urban index of a site, while the diversity of piscivores and carnivores increased with surrounding water coverage within 100 m. Mangrove specialist diversity decreased logarithmically with the extent of surrounding urban coverage. There were no detected relationships linking avian or anuran diversity with the diversity of surrounding vegetation classes, or with site floral composition or structure.

**Figure 3:**
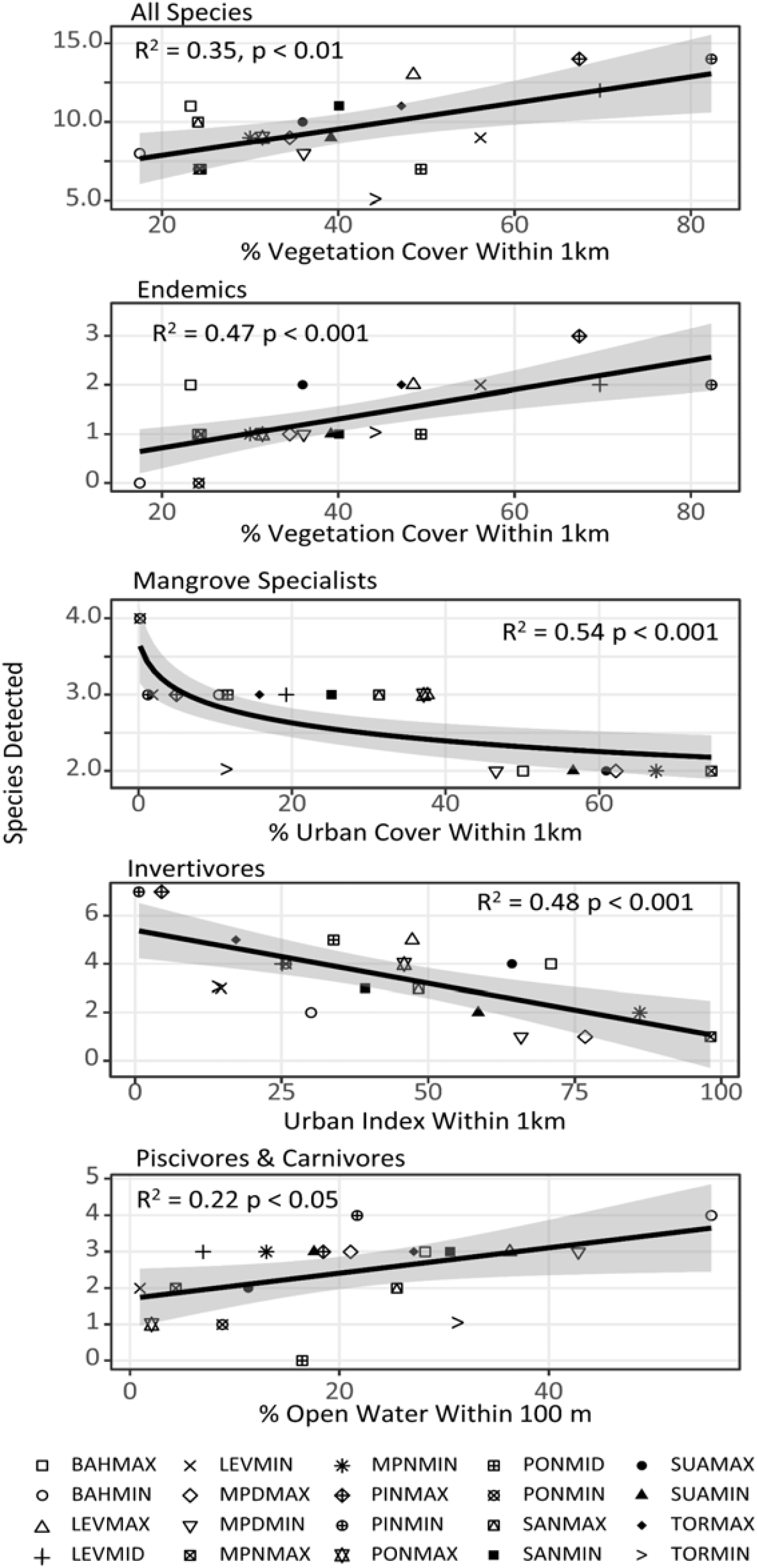
Diversity off all species, as well as a number of groups of species, were strongly correlated with surrounding land cover. Shaded areas are standard errors of the linear models.

The influence of urbanization on the faunal community was also evident in non-metric dimensional scaling (Figure 4). Separation of sites by their species’ detections resulted in some clustering of the most urban sites, as well as some separation from the least urban sites, but with a stress value of 0.16, the ordination should be interpreted with some caution. The most urban sites were mostly associated with the presence of both *Eleutherodactylus species*, as well as *Megaceryle alcycon, Coereba flaveola, Zenaida spp*., and *Seiurus noveboracensis*. Species associations for the least urban sites could not be determined due to a relative lack of clustering compared to the most urban sites, suggesting these sites are less similar to each other than are the urban sits. Further evidence for species specific associations with urbanization are presented in individual linear models.

**Figure 4:**
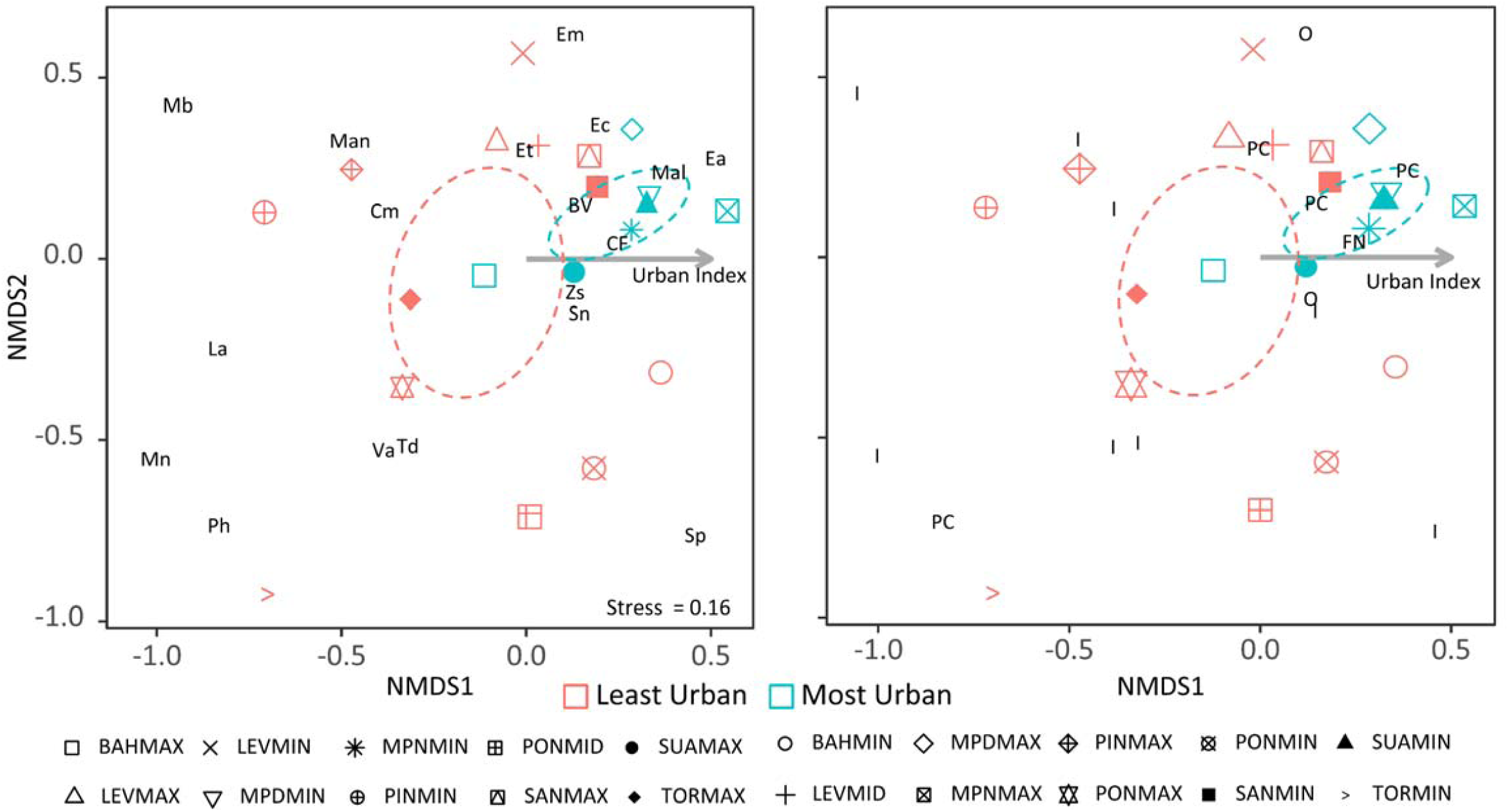
Non-metric dimensional scaling of all sites based on the detection frequencies of their species. Species associations are shown on the left, with the first letter(s) of their genus and species names, and dietary guilds are shown on the right, as abbreviated from Table 1. An overlay vector of the urban index shows in which general direction urbanization increases, and the ordinations are rotated so that this vector is horizontal for easier interpretation. Sites are colored as those with an urban index greater than the median (most urban) and those with an index value less than the median (least urban). The most urban sites are more clustered than the least urban sites, suggesting the former are more similar to each other.

The detection frequency of several species was also strongly correlated with surrounding land cover and urbanization metrics (Figure 5). Detections of both *Eleutherodactylus* species, as well as *Coereba flaveola* increased as populations density increased. Detections of *Megaceryle alcyon* also increased with urbanization, this time represented by the percent of surrounding urban coverage. *Coccyzus minor* showed the opposite trend, with detections decreasing as the surrounding urban index increased. Finally, detections of *Seirus noveboracensis* decreased with surrounding vegetation cover, although there was no trend with any of the urban metrics.

**Figure 5:**
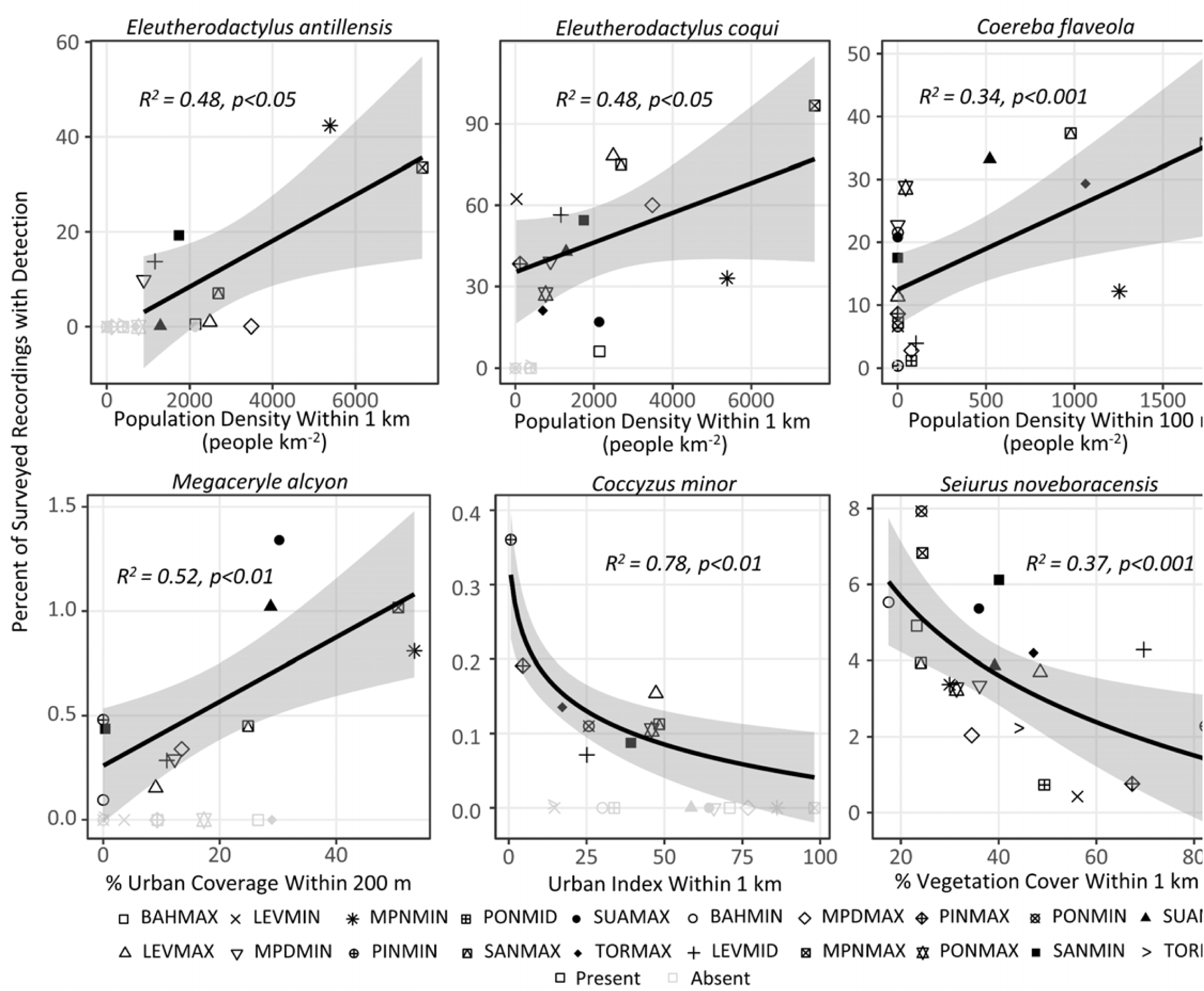
Individual species detection frequencies with surrounding land cover and urbanization metrics. Potential synanthropic species are indicated by an increase in detection frequencies with urbanization, while misanthropes are indicting by declining detections with increasing urbanization.

Soundscape analysis showed a significant relationship between ASU at a site and that site’s diversity (Figure 6). This relationship was distinct for sites within 0.5 km of low flying commercial airplanes, and those that were not. For those that were not influenced by the aircraft, the diversity ∼ ASU model predicted a linear increase of roughly one species for every 2.3% increase in the ASU. For those within the airport flight path, this relationship was one species for every 5.9% increase in ASU.

**Figure 6:**
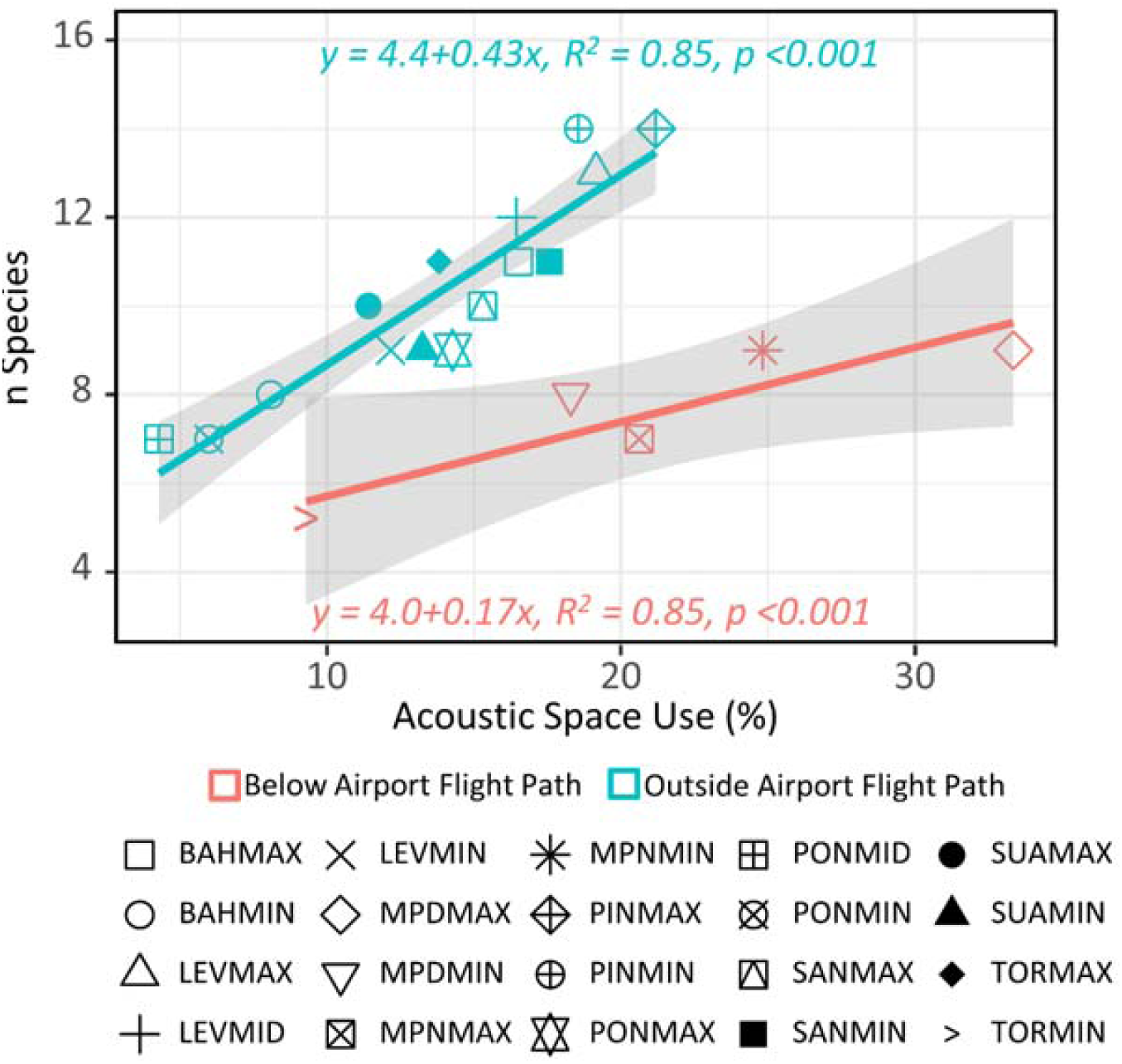
ASU is a strong predictor of site diversity, and can be used as a relatively rapid alternative to individual species detections. This relationship is strongly dependent on the associated anthrophony, with nearby commercial airplanes having the greatest influence in these sites.

## Discussion

Previous studies on urban bird communities have consistently shown an influence of the urban landscape on community structure and diversity, which is largely attributed to the diversity and availability of surrounding habitats and resources. Studies in mangrove systems are much less abundant but show similar trends. This study found both corroborating and contradictory findings in comparison to other studies, but largely shows that both vegetation cover and urbanization are highly influential in structuring mangrove faunal communities. Further, because of the high proportion of invertivores in the mangroves of Puerto Rico, and based on previous findings on the role of insects in driving forest faunal communities, it is likely that insects and their diversity are also driving anuran and avian communities in Puerto Rico’s mangrove habitats. Based on the findings of this study, these faunal communities can continue to be monitored for relative changes in diversity using the relatively easily calculated ASU and its strong correlation with avian and anuran diversity at each site.

The mangrove avifauna diversity of twenty-four species was only two species shy of that found by Acevedo & Aide (2008), although with some differences in specific species. This study detected few of the migrants, as well as some of the resident species observed by Acevedo and Aide, but compensated in detecting more resident species than the later. Further, in contrast with Acevedo and Aide, this study found no significance of the floral community or flooding variables on the faunal community. This is probably because the surveys conducted in Acevedo and Aide were done so in relatively large forested areas, with little variation in surrounding landcover between sites, in comparison to the smaller forest fragments of this study, which was purposely conducted in the greatest possible variation in landcover. Thus, when a large variation in landcover is present between sites, it will dominate the influence on faunal communities, but when surrounding landcover is more similar between sites, the influence of floral composition and hydrology will be more important.

Also unlike Acevedo and Aide (2008), the current study found a significant difference in native species presence between March and November of 2017. Further, detections of *Coereba flaveola,* a common native throughout the island, dropped across all sites between March and November. While seasonal variations in climate or breeding cycles may account for this difference, it is likely the passage of hurricane María over the island also had some influence. This disturbance resulted in widespread defoliation, which may have disproportionately affected this species by removing all flowers and fruits, its primary food source.

In this study, surrounding vegetation and urban coverage were the strongest predictors of mangrove faunal acoustic diversity. Both overall species and endemic species diversity were best predicted to increase with surrounding vegetation cover within one km. Although surrounding mangrove coverage was also a significant positive predictor for diversity, vegetation in general was strongest. This suggests that any vegetation type, not just mangroves, will support high faunal diversity in these forests. Mangrove specialist bird species, however, declined logarithmically with surrounding urban coverage, which was corroborated by declining detection frequencies of *Coccyzus minor* with increasing urban index. This suggests these species are more sensitive to urbanization than non-specialist species. This is likely because as urbanization increases and mangrove coverage is lost, replacement vegetation is not suitable habitat for specialists, which must move on to larger mangrove forests. Other studies have suggested mangrove specialists are capable of inhabiting urban mangroves (Lim and Sodhi, 2004; Noske, 1995; Ward, 1968), but these were nectarivores, which are absent in Puerto Rico’s mangrove specialists. The only two mangrove specialists present in the most urban mangroves of this study were *Zenaida spp*, an omnivore, and *Seirurs noveboracensis*, an invertivore. Its possible the *Zenaida* observed in these sites was the non-specialist *Zenaida asiatica*, leaving *Seirus noveboracensis* as the only urban mangrove specialist in Puerto Rico. As an invertivore, this species might be predicted to decline in urban areas, as did most of the others, but it instead responded positively to declining vegetation. It is unclear what drives this relationships, but this species in known only to roost in mangroves and move to foraging grounds during the day (Smith et al., 2008). Thus, mangroves with less surrounding vegetation, regardless of urbanization, may provide optimal foraging grounds.

Decreases in insectivorous mangrove avifauna have also been noted in another urban mangrove system (Lim and Sodhi, 2004), but studies in terrestrial systems suggest urban landscapes both favor and discourage use by avian insectivores (Chace and Walsh, 2006; Lancaster and Rees, 1979). Its unclear what drives these contradictions across systems, but in Puerto Rico urbanization disproportionately drives down the diversity of invertivores, leaving frugivores/nectarivores, omnivores and some 32 33 piscivores/carnivores more associated with urban habitats. The one frugivore/nectarivore in this study, *Coereba flaveola*, is likely drawn to urban forests due to their proximity to residential gardens, which harbor a greater variety of fruit and nectar produced by native and non-native plants (Parsons et al., 2003; Sewell and Catterall, 1998). As for piscivores/carnivores, Ardeidae were mostly associated with urban forests and these species have previously been noted in a variety of urban wetlands, suggesting relatively high synanthropy within this family (Fidalgo-De Souza, 2009; Samatha et al., n.d.; Scherer et al., 2006). Another urban associated piscivore, *Megaceryle alcyon*, was noted in the same sites in San Juan from over one-hundred years ago (Evermann et al., 1902), and its detection frequency increased as urban coverage increased, suggesting the species is also a synanthrop. The other piscivores/carnivores are the raptors, which have been generalized as being relatively well-suited to urban landscapes (Chace and Walsh, 2006). However, *Megascops nudipes* and *Pandion haliaetus*, were both associated with the least urban areas of the study, the former of which was relatively rare and only detected in well forested areas, and the later of which was associated with expansive open water. Only one invertivore, *Seiurus noveboracensis*, was associated with urban landscapes, likely due to the its previously mentioned affinity for sparse vegetation. However, one possible alternative explanation to the patterns seen in *Seirus noveboracensis*, as well as in *Megaceryle alcyon* and *Coccyzus minor*, is that the change in habitat availability in the least and most urban sites results in decreases and increases of population density, respectively, as more individuals are forced into small urban mangrove forests, resulting in a higher detection frequency compared to more expansive forests where populations are less dense. In any case, their use of these spaces suggests a relatively high suitability for urban mangrove habitats.

The three anurans of this study also showed some tendency towards either synanthropic or misanthropic habitat preferences. Both members of *Eleutherodactylus* showed increasing detection frequency with population density, but *Leptodactylus albilabris* was absent from the most urban sites. This is likely because the former exhibit direct development and have no tadpole stage that requires pooled water (Townsend and Stewart, 1985), while *Leptodactylus albilabris* does (Dent, 1956). Thus, *Eleutherodactylus spp*. are able to utilize urban mangrove habitats more so than is *Leptodactylus albilabris* probably by avoiding contact with surface waters and instead maintaining water balance through freshwater from foliage and the atmosphere (Taigen et al., 1984). Although previous studies have noted the presence of *Eleutherodactylus spp.* in urban landscapes, none have suggested a preference for urban areas. The high frequency of detections in the urban mangroves in this study may simply be a result of limited vegetated habitat in urban areas, making the mangroves and the estuarine-terrestrial interface the most accommodating space in these landscapes.

Soundscape analysis and ASU provide an opportunity for facilitating the continued acoustic monitoring of the mangroves of Puerto Rico. By understanding how a forest’s soundscape is influenced by both the species diversity and the surrounding anthrophony, it’s possible to conduct rapid diversity surveys without the time and labor demanding process of creating species specific acoustic models. This study found a strong relationship between a site’s ASU and the diversity of noisy species. It also found how a persistent anthropogenic sound source, low flying commercial airplanes, influence the relationship between ASU and species diversity. Further, 80% of the species were detected in the first 500 minutes of recording across all twenty sites, and extrapolations predict a 15 hectare sampling area should yield roughly twenty-five to thirty species in each watershed. This information can be used to predict the same diversity in future surveys, which can help monitor the faunal communities of Puerto Rico’s mangroves.

As the Caribbean has been identified as one of five biodiversity hotspots predicted to undergo relatively rapid urbanization in the twenty-first century (Seto et al., 2012), a rise of urban mangroves there may bring changes to faunal communities. This study showed distinct faunal communities between the most and least urban mangroves of Puerto Rico, with the former being less diverse and more homogenous than the latter. Although the most urban mangroves offer habitat for some of the synanthropic species, other species seem to be avoiding these forests, and will be the most vulnerable to ongoing coastal urbanization. Thus, conserving large expansive mangrove forests with minimal urban influences will help ensure avian and anuran diversity, especially of endemics and mangrove specialists. At the same time, highly urban forests can be managed to optimize cultural ecosystem services that take advantage of both avian and anuran use of these habitats. Both strategies will be important for the long-term sustainability of these social-ecological systems.

## Acknowledgments

Funds were provided by the United State’s Department of Agriculture’s Forest Service International Institute of Tropical Forestry. Dr. Marconi Campos Cerqueira provided species identification confirmations. Ariel Lugo reviewed preliminary drafts.

